# PyMethylProcess - highly parallelized preprocessing for DNA methylation array data

**DOI:** 10.1101/604496

**Authors:** Joshua J. Levy, Alexander J. Titus, Lucas A. Salas, Brock C. Christensen

## Abstract

**Summary:** The ability to perform high-throughput preprocessing of methylation array data is essential in large scale methylation studies. While R is a convenient language for methylation analyses, performing highly parallelized preprocessing using Python can accelerate data preparation for downstream methylation analyses, including large scale production-ready machine learning pipelines. Here, we present a methylation data preprocessing pipeline called PyMethylProcess that is highly reproducible, scalable, and that can be quickly set-up and deployed through Docker and PIP.

**Availability and Implementation:** Project Name: PyMethylProcess

Project Home Page: https://github.com/Christensen-Lab-Dartmouth/PyMethylProcess. Available on PyPI as *pymethylprocess*.

Available on DockerHub via *joshualevy44/pymethylprocess*.

Help Documentation: https://christensen-lab-dartmouth.github.io/PyMethylProcess/

Operating Systems: Linux, MacOS, Windows (Docker)

Programming Language: Python, R

Other Requirements: Python 3.6, R 3.5.1, Docker (optional) License: MIT

## 1. Implementation

DNA methylation plays a critical role in cell fate determination during development and regulation of transcription throughout life (Ji *et al*., 2010; Khavari *et al*., 2010; Rönnerblad *et al*., 2014). Studies that measure DNA methylation in large numbers of human biospecimens typically use the Illumina Infinium BeadArray platforms known as 27K, 450K and 850K/EPIC arrays (Bibikova *et al*., 2011, 2009; Moran *et al*., 2016). Popular R packages used for DNA methylation data processing and analysis include *minfi* (Aryee *et al*., 2014)*, ENmix* (Xu *et al*., 2016), or *meffil* (Min *et al*., 2018). However, a straightforward and tractable approach to perform data quality control and functional normalization in bulk to prepare the data for use in the object-oriented environment is lacking. Here, we introduce a simple and easy-to-use command line interface that makes methylation analyses more object oriented and ready for use in down-stream analyses such as immune cell proportion estimation, and python-oriented Epigenome-Wide Association Studies (EWAS) studies. In addition to more traditional methylation-based analyses, popular machine learning libraries such as scikit-learn, keras, and tensorflow (Abadi *et al*., 2016; Pedregosa *et al*., 2011) become more accessible to both machine learning researchers without an epigenetic background and epigenetic researchers with a limited machine learning background.

PyMethylProcess is a pip-installable command line interface built using Python 3.6 that interfaces with *minfi, ENmix*, and *meffil* in R via rpy2 (Gautier, 2010) to allow users to preprocess and set-up their methylation array data for downstream analyses included machine learning, presenting unique methylation datatypes that are built for the use of python classification, clustering, dimensionality reduction and regression algorithms such as UMAP, random forest, neural networks, k-nearest neighbors, and HDBSCAN (Campello *et al*., 2013; Ho, 1995; McInnes *et al*., 2018). Eight Python classes have been introduced to handle the following tasks: package installation (*PackageInstaller* installs any R/bioconductor package via python), data acquisition from TCGA and GEO (*TCGADownloader*) and formatting (*PreProcessPhenoData*), parallelized quality control (QC), principal component selection via kneedle (Satopaa *et al*., 2011), and raw, quantile, noob, and functional normalization using *minfi, meffil, ENmix* (*PreprocessIDAT*), imputation (*ImputerObject)*, feature selection, final preparation and storage for machine learning applications (*MethylationArray[s]* store beta and phenotype data), and a basic machine learning API class *MachineLearning* that trains any scikit-learn-like model on *MethylationArray* objects. These datatypes are used to handle complex preprocessing calculations, while being abstracted away via a convenient command-line interface. Additional commands, such as the removal of non-autosomal sites, SNP removal (either via QC methods or post-QC by subsetting CpGs that are not in a list of CpGs supplied by *meffil* for the respective array platform), and reference-based cell-type estimation (constrained projection/quadratic programming) (Houseman *et al*., 2012; Jaffe and Irizarry, 2014; Salas *et al*., 2018), and class methods are available in the help documentation. In addition, a visualization module generates interactive 3-D representations of the data using UMAP and Plotly (Modern Analytic Apps for the Enterprise) for further inspection. The beta values and phenotype data from the *MethylationArray* objects can easily be written to csv format using *write_csv*, and the command line interface can speed up the time it takes for researchers to set-up their standardized methylation data for downstream analyses.

**Figure 1.**
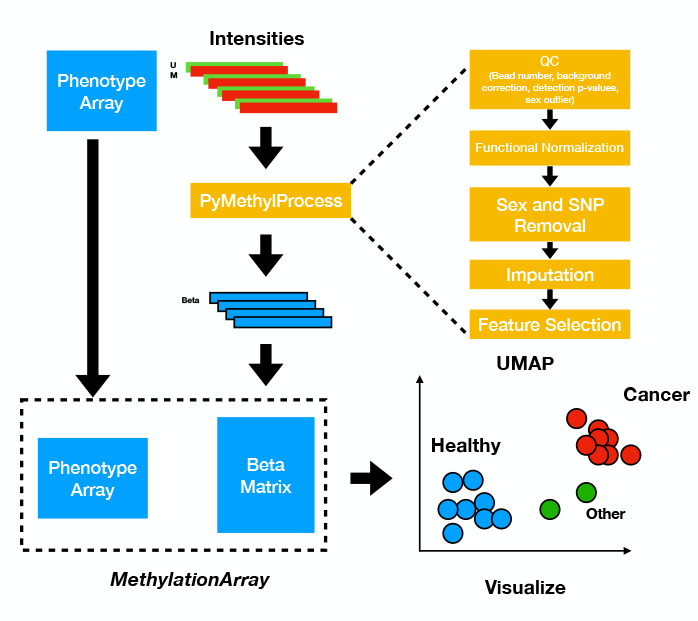
Flow Diagram for *PyMethylProcess*.

The pipeline differs from other python frameworks such as pyMAP (Mahpour, 2016) and GLINT (Rahmani *et al*., 2017) in functionality. pyMAP only operates on the 450k framework, relying on user specific csv annotation files and preprocessed Genome Studio txt files as its input. pyMAP only performs graphical exploration for candidate CpGs, CpG Island feature subsetting, and export to BED files for downstream analyses. Similarly, GLINT requires a txt phenotype file and either a preprocessed beta values txt files or a R data.frame methylation object (in a RData file) as its inputs. GLINT stores beta values and covariate information as a binary “glint” file. GLINT was designed for EWAS analysis, including reference-based and reference-free estimations, imputed genetic structure, and statistical models (linear, logistic and linear mixed effects models). However, it relies on preprocessed data, with some limited quality control options and as so it could benefit from preprocessed data generated in our pipeline. In addition, GLINT is not designed to export this information to perform user customized downstream machine learning analyses.

## 2. Results

Here we demonstrate some of *PyMethylProcess*’s capabilities, performing preprocessing of seven different datasets (GSE87571, GSE81961, GSE69138, GSE42861, GSE112179, GSE109381, TCGA Pan-cancer)(Capper *et al*., 2018; Johansson *et al*., 2013; Li Yim *et al*., 2016; Pai *et al*., 2018; Pidsley *et al*., 2013, 2013; Salas *et al*., 2017; Soriano-Tárraga *et al*., 2018) from the HumanMethylation450 BeadArray (450K array), and the HumanMethylationEPIC BeadArray (850K array), platform, derived from GEO, most of which benchmarked for loading, QC and normalization time. After preprocessing, each of the below data sets were split into 70% training, 10% validation, and 20% test sets for downstream prediction.

**Table 1.**
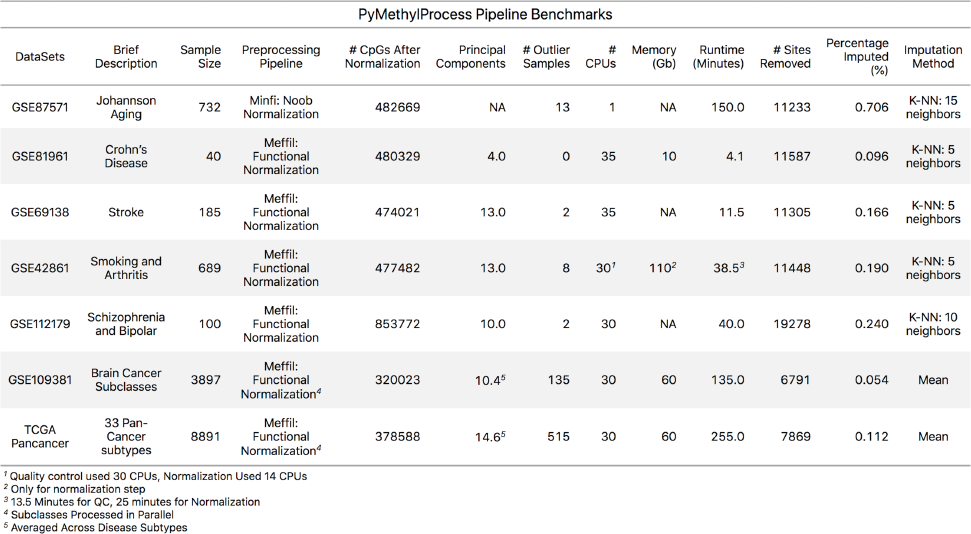
Benchmark and Preprocessing Results

## 3. Benefits and Future Direction

The *PyMethylProcess* streamlines the process of preprocessing Methylation Array data while making Methylation data highly accessible and standardized for the Python machine learning community. *PyMethylProcess* is open source software, and additional development based on the community needs may be added. The authors of this paper would like to solicit issues and pull requests from the greater bioinformatics community on GitHub.

Future development will expand functionality to other preprocessing pipelines such as Watermelon and BigMelon (Gorrie-Stone *et al*.) and feature importance evaluations using measures such as the Gini index will be included. In addition, this command-line tool is available for use via Docker (*joshualevy44/pymethylprocess*) (Boettiger, 2015), making this analysis easily sharable, standardized, and operating system agnostic. The entire workflow will be wrapped using Common Workflow Language (CWL) (Amstutz *et al*., 2016), making the entire analysis executable at the click of a button, highly reproducible, and easily sharable.

*PyMethylProcess* is installable from PyPI via the name *pymethylprocess*, and source code ***can be found on GitHub at: https://github.com/Christensen-Lab-Dartmouth/PyMethylProcess***

## Acknowledgements

JJL conceived of idea and implementation, programmed and tested pipeline on datasets, wrote text; AJT tested pipeline; LAS provided technical support to streamline and debug important aspects of the pipeline; BCC provided research direction support; all authors participated in editing the text.

## List of Abbreviations

EWAS: Epigenome-Wide Association Studies
QC: Quality Control
CWL: Common Workflow Language
450K: HumanMethylation450 BeadArray
850K: HumanMethylationEPIC BeadArray

